# Concurrent Atomic Force Spectroscopy

**DOI:** 10.1101/293506

**Authors:** Carolina Pimenta-Lopes, Carmen Suay-Corredera, Diana Velázquez-Carreras, David Sánchez-Ortiz, Jorge Alegre-Cebollada

**Author notes:** Correspondence to Jorge Alegre-Cebollada. equal contribution (Carolina Pimenta-Lopes, Carmen Suay-Corredera). Department of Physiological Sciences, University of Barcelona, Spain (Carolina Pimenta-Lopes).

## Abstract

Force-spectroscopy by Atomic Force Microscopy (AFM) is the technique of choice to measure mechanical properties of molecules, cells, tissues and materials at the nano and micro scales. However, unavoidable calibration errors of AFM probes make it cumbersome to quantify modulation of mechanics. Here, we show that concurrent AFM force measurements enable relative mechanical characterization with an accuracy that is independent of calibration uncertainty, even when averaging data from multiple, independent experiments. Compared to traditional AFM, we estimate that concurrent strategies can measure differences in protein mechanical unfolding forces with a 6-fold improvement in accuracy and a 30-fold increase in throughput. Prompted by our results, we demonstrate widely applicable orthogonal fingerprinting strategies for concurrent single-molecule nanomechanical profiling of proteins.

## INTRODUCTION

The development of Atomic Force Microscopy (AFM) has enabled imaging the nanoscale with unprecedented length resolution, revolutionizing nanotechnology, materials science, chemistry and biology ^1-3^. AFM is based on the detection of interaction forces between a sample and a microfabricated cantilever, the force probe of the technique. Traditionally, low-scanning speeds have limited the reach of AFM; however, recent developments involving miniaturized cantilevers have achieved imaging at video frame rates, launching the field of high-speed AFM^4^. The ability of AFM to measure forces at the nano and micro scales can be exploited to quantify mechanical parameters such as stiffness, viscoelasticity, or adhesion forces, while simultaneously imaging surface topology ^2,5,6^. Due to its force sensitivity down to picoNewtons (pN), AFM-based force spectroscopy, or simply atomic force spectroscopy, can also be used to examine single-molecule dynamics and ligand-receptor interactions under force ^7-9^, of relevance for cellular stiffness, mechanosensing and mechanotransduction ^10-13^.

A key limitation of atomic force spectroscopy stems from uncertain calibration of the spring constant (*k_sc_*) of the AFM cantilever, which is needed to determine force values ^14^. Different calibration methods to estimate *k_sc_* have been developed, differing in their simplicity, damage to the cantilever tip, experimental compatibility and associated uncertainty. Estimates of calibration uncertainty up to 25% are usually reported ^15^; however, even higher inaccuracies can result from defects in individual cantilevers, and from unpredictable changes in *k_sc_* during experimentation due to material deposition, mechanical drift and wear ^7,16^.

Force calibration uncertainties in AFM lead to inaccurate quantification of mechanical properties, a situation that is worsened in modes where the elusive geometry of the cantilever tip is required to extract mechanical information ^17^. Indeed, relative AFM studies to characterize mechanical modulation of proteins, materials and cells are challenging, since they necessitate multiple experiments, each one affected by different calibration errors ^18,19^. As a result, there is a pressing need to develop methods that can overcome inaccurate AFM force calibration ^18^. The traditional approach to increase statistical power is to repeat experiments, since individual calibration errors are more probable to be averaged out as more experiments are included in the analysis ^20^. The drawback is a considerable loss of throughput of the technique.

In theory, a manner to overcome calibration errors in relative atomic force spectroscopy is to measure the samples concurrently in a single experiment, using one cantilever under the same calibration parameters ^21^. Indeed, mechanical characterization of several proteins in a single experiment has been achieved using microfluidics, on-chip protein expression and force-spectroscopy measurements in custom-built atomic force/total internal reflection fluorescence microscopes ^22,23^. However, this advanced technology is not available to most AFM users, and the extent of improvement in performance by concurrent AFM remains unexplored.

Here, we use error propagation analysis and Monte Carlo simulations to examine how force calibration errors impact determination of mechanical properties of proteins by atomic force spectroscopy, and demonstrate the remarkable improvement that stems from concurrent measurements. Unexpectedly, we find that averaging results from multiple concurrent experiments retains statistical power, leading to drastically improved accuracy and throughput in the determination of relative mechanical properties by AFM. Prompted by our findings, we have developed orthogonal fingerprinting (OFP) as a simple and widely applicable strategy for concurrent single-molecule nanomechanical profiling of proteins.

## RESULTS

### Interexperimental variation of mechanical properties obtained by traditional atomic force spectroscopy

To understand how errors in calibration of atomic force microscopes lead to inaccurate determination of mechanical properties, we have measured protein mechanical stability by single-molecule force-spectroscopy AFM ^7^ (Figure 1A, Supplementary Figure 1). This AFM mode is very well suited to examine propagation of calibration errors since protein unfolding forces are obtained directly from experimental data and do not depend on further modelling or approximations.

**Figure 1.**
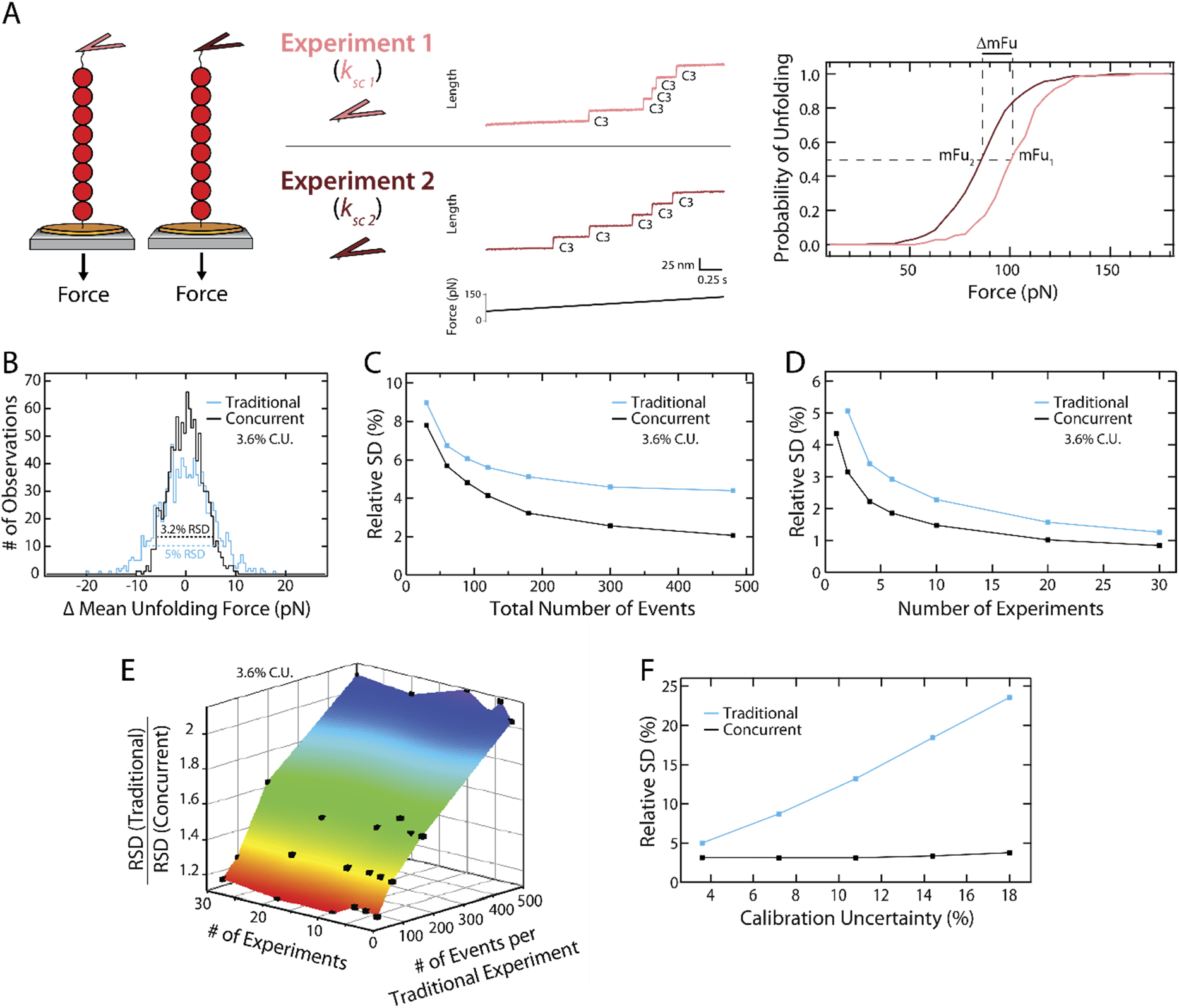
Concurrent mechanical profiling by atomic force spectroscopy circumvents inaccurate force calibration. **(A)** *Left:* Schematic representation of the traditional strategy used to measure mechanical stability of proteins by single-molecule atomic force spectroscopy, in which data are obtained in multiple, independent experiments under different force calibration parameters (see also Supplementary Figure 1). *Middle*: Results from two independent force-spectroscopy AFM experiments in which a (C3)_8_ polyprotein is pulled under a 40 pN/s linear increase in force. Due to uncertain calibration of the cantilever’s spring constant (*k_sc_*), force values are affected by errors that differ between experiments. We show two individual unfolding traces of the (C3)_8_ polyprotein, in which mechanical unfolding events of individual C3 domains are detected by increases of 24 nm in the length of the polyprotein. *Right*: Experimental cumulative unfolding probability distributions obtained from 117 (Experiment 1) and 191 (Experiment 2) C3 unfolding events. The corresponding *mF_u_* values are 98.7 and 82.9 pN, respectively. **(B)** Distributions of *∆mF_u_* obtained by Monte Carlo simulations, considering the same total number of experiments and unfolding events for both traditional (blue) and concurrent measurements (black). We considered 2 experiments, 200 total unfolding events per protein, and a 3.6% force calibration uncertainty (C.U.) **(C)** Keeping the number of experiments constant (n_exp_=2), the Relative Standard Deviation (RSD) of the distribution of *∆mF_u_* decreases with the total number of unfolding events both in traditional (blue) and in concurrent measurements (black). **(D)** Keeping the same number of events per experiment (n_events per experiment_ = 200, in the traditional approach, and 100 in the concurrent strategy), the RSD of the distribution of *∆mF_u_* decreases with the number of experiments (blue: traditional strategy; black: concurrent measurements). **(E)** The relative improvement in the RSD of *∆mF_u_* distributions obtained by the concurrent strategy increases with the number of events per experiment, and remains fairly insensitive to the number of averaged experiments. **(F)** RSD of the distributions of *∆mF_u_* at increasing calibration uncertainties for the traditional (blue) and the concurrent (black) strategies. The remaining simulation parameters are the same as in panel B. In panels B-F, the number of events per experiment and protein in the concurrent approach was half of the number of events in the traditional strategy so that RSDs are compared between conditions with equal total number of events.

We first measured the resistance to mechanical unfolding of the same protein in different, independently calibrated atomic force spectroscopy experiments. We produced a polyprotein containing eight repetitions of the C3 domain of human cardiac myosin-binding protein C (Supplementary Figure 2, Supplementary Text 1) and subjected individual (C3)_8_ polyproteins to a linear increase in force of 40 pN/s using a force-clamp atomic force microscope. Results from two independent experiments are shown in Figure 1A. Mechanical force triggers the stochastic unfolding of individual C3 domains. These unfolding events are detected as 24 nm step increases in the length of the polyprotein (Figure 1A, *middle*; Supplementary Figure 3A). We measured the force at which the unfolding events occur and calculated distributions of unfolding forces ^24^. Despite the fact that both distributions are well defined (n > 115 events), the difference in their mean unfolding force (*∆mF_u_*) is 19% (Figure 1A, right). The magnitude of this interexperimental variation is in agreement with the spread of *mF_u_* values reported in the literature for other proteins ^20^. These differences can mask comparable changes in mechanical properties induced by disease-causing mutations ^19^ or posttranslational modifications ^25^, which is a key limitation of traditional single-molecule atomic force spectroscopy.

Interexperimental variations in *mF_u_* of proteins are typically interpreted in terms of different errors in the calibration of AFM cantilevers, a procedure that can entail 25% uncertainty ^7,14,15,22^. We used Monte Carlo simulations to examine how errors originating from uncertain cantilever calibration propagate to *mF_u_* values measured by atomic force spectroscopy experiments (see Methods). Briefly, in each simulated experiment we impose an error to the force that is randomly drawn from a normal distribution defined by a Relative Standard Deviation (RSD) that is equal to the considered calibration uncertainty. After definition of the error in force for a cycle, a kinetic Monte Carlo algorithm is used to obtain distributions of unfolding forces for a given number of protein domains subjected to a nominal 40 pN/s increase in force. For each one of the 1,000 cycles of the simulation, we calculate the corresponding *∆mF_u_* considering a certain number of independent experiments, each one affected by a different calibration error. Simulations return the distribution of *∆mF_u_* values obtained in the 1,000 cycles. The spread of the distribution, which is a measure of the accuracy of atomic force spectroscopy data, is quantified by its RSD.

Using our Monte Carlo procedure, we have simulated mechanical protein unfolding under a modest 3.6% force calibration uncertainty. This value is a good estimate of the lowest uncertainty that can be achieved by the thermal fluctuations method, typically used in single-molecule AFM studies (Supplementary Text 2, Supplementary Figure 4) ^7^. Figure 1B shows the simulated distribution of *∆mF_u_* obtained from two independent experiments in which the same protein is probed (200 unfolding events per experiment). It is remarkable that two mean unfolding forces obtained in different cycles can differ by more than 25% (Supplementary Figure 5A). Hence, although conservative, a mere 3.6% inaccuracy in cantilever calibration can explain considerably higher differences in *mF_u_* obtained in traditional atomic force spectroscopy experiments.

### Accuracy in *∆mF_u_* obtained by concurrent atomic force spectroscopy is insensitive to calibration uncertainty

We have used our Monte Carlo simulations to estimate the accuracy in *∆mF_u_* achieved by concurrent measurement of the mechanical stability of two proteins. Considering equal total number of events and experiments, and a 3.6% calibration uncertainty, we find that the RSD of the distribution of *∆mF_u_* decreases from 5.0% to 3.2% when measurements are taken concurrently (Figure 1B). RSD values are further reduced at higher number of unfolding events, as expected from better definition of the distribution of unfolding forces (Figure 1C, Supplementary Figure 5B). Unexpectedly, averaging multiple concurrent experiments leads to reductions in RSD, despite the fact that each individual experiment is performed under different calibration parameters (Figure 1D). Increasing the number of events or experiments also results in lower RSD values when proteins are probed in traditional, separate experiments (Figure 1 C,D). We find that the relative improvement in RSD achieved by concurrent over traditional atomic force spectroscopy increases with the number of unfolding events per experiment, and remains fairly constant at increasing number of experiments (Figure 1E). Hence, we conclude that averaging independent atomic force spectroscopy experiments in which two proteins are probed concurrently retains statistical power, even if those individual experiments are affected by different calibration errors.

All our simulations in Figure 1B-E were run considering 3.6% uncertainty in force calibration, which is much smaller than usually reported ^7,14,15,22^. Hence, we estimated the RSD of the distribution of *∆mF_u_* at increasing calibration uncertainties. As expected, higher calibration uncertainties lead to much increased RSD in traditional atomic force spectroscopy, whereas the RSD of concurrent distributions remains insensitive to calibration uncertainty (Figure 1F), even when data from several independent experiments are averaged (Supplementary Figure 6A).

### Orthogonal fingerprinting enables concurrent mechanical characterization of proteins by atomic force spectroscopy

Our simulation results show that under a modest 3.6% uncertainty in force, concurrent atomic force spectroscopy measurements can reach the same level of accuracy in *∆mF_u_* with 2-4 times less experiments than the traditional approach (Figure 1D). Furthermore, at high values of calibration uncertainty, the accuracy in *∆mF_u_* of concurrent measurements can be 6 times higher than in the traditional approach (Figure 1F). These remarkable improvements in throughput and accuracy prompted us to design a general strategy for concurrent measurements of mechanical properties of proteins that can be readily applied to any AFM-based force spectrometer.

Having methods to identify single-molecule events is a fundamental requirement of force-spectroscopy AFM. In the case of mechanical characterization of proteins, this need is fulfilled by the use of polyproteins, which provide molecular fingerprints that easily discriminate single-molecule events from spurious, non-specific interactions ^26,27^ (Supplementary Figure 3). As exemplified in Figure 1A, mechanical unfolding of polyproteins produce repetitive events whose length fingerprints the domain of interest. If two polyproteins are to be measured concurrently in the same experiment, it is imperative that they have different fingerprinting unfolding lengths. Here, we propose a widely applicable manner of achieving such orthogonal fingerprinting (OFP) through the use of heteropolyproteins, in which marker proteins are fused to the proteins of interest ^28^. Since OFP identifies proteins through the unfolding length of the marker domains, proteins of interest to be compared can have the same unfolding length (e.g. mutant or biochemically modified proteins). To test whether heteropolyproteins can be employed to achieve OFP during concurrent measurement of proteins by atomic force spectroscopy, we have measured the mechanical stability of the C3 domain in different polyprotein contexts.

We first followed a single-marker strategy using the heteropolyprotein (C3-L)_4_ (Figure 2, Supplementary Figure 2 and Supplementary Text 1). In (C3-L)_4_, we used protein L as a marker since its unfolding length is different from the one of C3 ^20^. Indeed, mechanical unfolding of (C3-L)_4_ under a 40 pN/s ramp results in the appearance of 16 and 24 nm steps, which correspond, respectively, to the unfolding of L and C3 domains (Figure 2B, *left*, Supplementary Figure 3B). We selected unfolding traces of (C3)_8_ and (C3-L)_4_, obtained in independent, traditional fingerprinting (TFP) experiments, and classified them according to the number of 16 and 24 nm events they contain. Our results show that a gating criterion of n(16nm) = 0 and n(24nm) > 2 unambiguously identifies traces coming from (C3)_8_, whereas traces resulting from (C3-L)_4_ can be safely assigned when n(16nm) > 1 and 0 < n(24nm) < 5 (Figure 2B, *right*). We analyzed 17 such TFP experiments and obtained distributions of unfolding forces for C3 in the context of both polyproteins, which we found to be very similar (*mF_u_* = 90. 7 and 88.4 pN for the homo and the heteropolyproteins, respectively, Figure 2C, Table). We used our Monte Carlo simulations to estimate the RSD associated to *∆mF_u_* that is expected from the actual number of experiments and events obtained (Figure 2C, Table, Supplementary Figure 7A).

**Figure 2.**
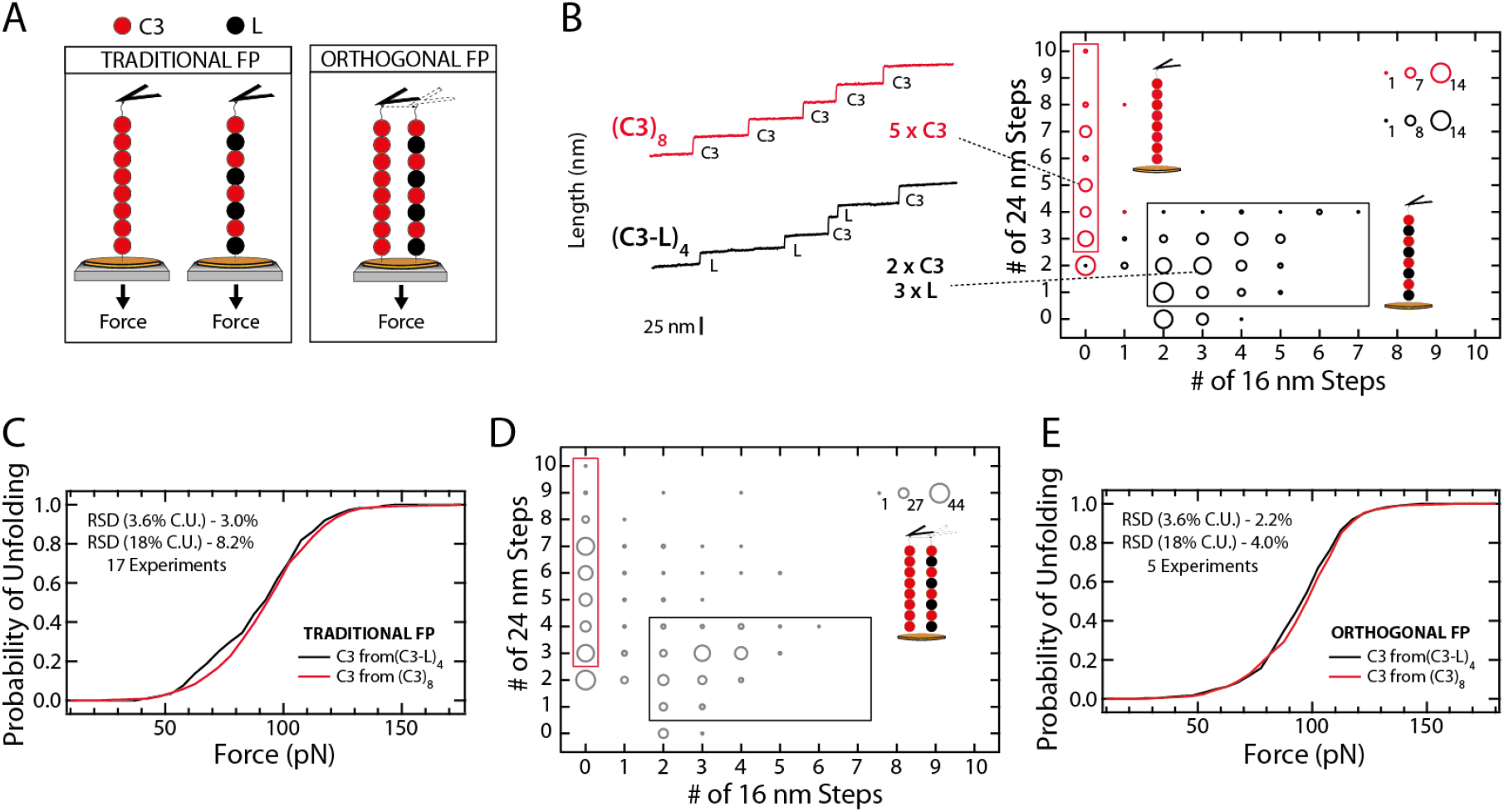
Orthogonal fingerprinting enables concurrent measurement of proteins by atomic force spectroscopy. **(A)** Schematic representation of the traditional and orthogonal fingerprinting (TFP and OFP, respectively) strategies. *Left*: in TFP, (C3)_8_ and (C3-L)_4_ are measured in different atomic force spectroscopy experiments. *Right*: In OFP both polyproteins are measured concurrently in the same experiment. **(B)** *Left*: representative unfolding traces of one (C3)_8_ molecule (red), in which five 24 nm unfolding events are detected, and one (C3-L)_4_ polyprotein (black) in which two 24 nm and three 16 nm unfolding events are observed. *Right*: Mechanical unfolding of (C3)_8_ (red) and (C3-L)_4_ (black) were measured in separate experiments. Individual traces were classified according to the number of 16 and 24 nm steps they contain. The plot shows the frequency of the traces that have different combinations of unfolding events, as indicated by the size of the dots. The rectangles represent gating strategies that unequivocally identify traces coming from (C3)_8_ or (C3-L)_4_. Number of traces analyzed: (C3)_8_, 59; (C3-L)_4_, 129. **(C)** Experimental cumulative unfolding probability distribution of the C3 domain in the context of (C3)_8_ (11 experiments, 1334 events, red) and (C3-L)_4_ (6 experiments, 177 events, black), following TFP. **(D)** (C3)_8_ and (C3-L)_4_ polyproteins were measured concurrently in the AFM. 468 traces were classified according to the number of 16 and 24 nm events they have. The plot shows the frequency of the traces that have different combinations of unfolding events, as indicated by the size of the dots. The gating criterion defined in panel B allows the classification of individual traces as resulting from (C3)_8_ (red rectangle) or (C3-L)_4_ (black rectangle). **(E)** Experimental cumulative unfolding probability distribution of the C3 domain in the context of (C3)_8_ (625 events, red) and (C3-L)_4_ (311 events, black), resulting from unfolding data obtained in 5 independent OFP experiments. RSD values in the insets of panels C and E are estimated using Monte Carlo simulations that consider extreme values of calibration uncertainty (C.U.) (see also Supplementary Figure 7A). The pulling rate in all experiments was 40 pN/s.

**Table.** Mechanical characterization of C3 by single-molecule atomic force spectroscopy. The table summarizes results regarding mechanical characterization of the C3 domain in the context of three different polyproteins, following traditional atomic force spectroscopy or OFP-based concurrent measurements. n_exp_ and n_events_ represent the number of experiments and unfolding events, respectively. RSDs of the distributions of ***∆mF_u_*** with respect to (C3-L)_4_ at two different calibration uncertainties (C.U.) were estimated by Monte Carlo simulations using the actual structure of the experimental datasets (Supplementary Table 1).

| AFM mode | Protein | $C3, mF_u$<br>(pN) | $n_{exp}$ | $n_{events}$ | RSD at 3.6%<br>C.U. (%) | RSD at 18%<br>C.U. (%) |
| --- | --- | --- | --- | --- | --- | --- |
| TFP | (C3)<sub>8</sub> | 90.7 | 11 | 1334 | 3.0 | 8.2 |
|  | (C3-L)<sub>4</sub> | 88.4 | 6 | 177 | n/a | n/a |
|  | (C3-Sumo1)<sub>4</sub> | 79.2 | 8 | 742 | 3.2 | 9.4 |
| OFP, single<br>marker | (C3)<sub>8</sub> | 96.3 | 5 | 625 | 2.2 | 4.0 |
|  | (C3-L)<sub>4</sub> | 93.4 | 5 | 311 | n/a | n/a |
| OFP, dual<br>marker | (C3-L)<sub>4</sub> | 90.0 | 14 | 873 | n/a | n/a |
|  | (C3-Sumo1)<sub>4</sub> | 89.5 | 14 | 1043 | 1.5 | 2.9 |

Following validation of the polyprotein gating criterion (Figure 2B), we measured (C3)_8_ and (C3-L)_4_ concurrently in OFP experiments. Single-molecule traces were classified according to the number of 16 and 24 nm steps they contain, and sorted as coming from the (C3)_8_ or (C3-L)_4_ before analysis of unfolding data (Figure 2D). As in the case of TFP experiments, the unfolding probability distributions of C3 in the context of (C3)_8_ and (C3-L)_4_ are very similar (*mF_u_* = 96.3 and 93.4 pN for the homo and the heteropolyproteins, respectively, Figure 2E, Table). However, only 5 OFP experiments were required to reach a lower RSD than in TFP, a 3 times higher speed of data acquisition (Figure 2C,E, Table, Supplementary Figure 7A).

### Dual-marker orthogonal fingerprinting overcomes confounding protein dimerization

In the atomic force spectroscopy experiments reported in Figures 1 and 2, polyproteins are picked up by the AFM cantilever through non-specific physisorption. Hence, experimental traces can contain different number of unfolding events (Figure 2B). Non-specific protein pickup also leads to the occasional appearance of traces containing more unfolding events than engineered domains in the polyprotein, an effect that results from polyprotein dimerization ^29^. For instance, in Figure 2B a few traces with n(24 nm) > 8 are identified when pulling from (C3)_8_. Comparison of Figures 2B and 2D identifies a new population of events at n(24nm) > 4 and n(16nm) > 1 that appear only when (C3)_8_ and (C3-L)_4_ are measured concurrently in the same experiment, which we interpret as heterodimers between (C3)_8_ and (C3-L)_4_. In the context of OFP, heterodimerization hampers proper assignment of traces, since there is a non-zero probability that some heterodimers are included in the gating region of (C3-L)_4_. As a result, a fraction of events coming from (C3)_8_ could be mistakenly assigned to (C3-L)_4_.

In general, the extent of protein dimerization in single-molecule AFM is dependent on the particular experimental conditions. Hence, heterodimerization poses a challenge to OFP, whose extent may vary depending on the system to study. However, we hypothesized that difficulties coming from protein dimerization could be overcome by using a second protein marker, since traces originating from dimers would be fingerprinted by the presence of both marker proteins. We chose the protein SUMO1 as a second marker because its unfolding length is different from those ones of C3 and protein L ^30^. We engineered the heteropolyprotein (C3-SUMO1)_4_ and pulled it in the AFM (Supplementary Figure 2). Two population of unfolding steps, at 20 nm and 24 nm are detected, corresponding to the unfolding of SUMO1 and C3, respectively (Supplementary Figure 3C).

Having two marker proteins enables gating criteria that are based exclusively on the presence of the marker domains, in a manner that protein dimers can be identified and excluded from the analysis (Figure 3A,B, Supplementary Figure 8). According to experiments in which (C3-L)_4_ and (C3-SUMO1)_4_ are measured separately, we used the gating criterion that n(16nm) > 1 and n(20nm) = 0 marks unfolding of (C3-L)_4_, and n(20nm) > 1 and n(16nm) = 0 defines unfolding of (C3-SUMO1)_4_, which only misclassifies 1 out of 136 traces. Following this gating criterion, we determined the distribution of unfolding forces of C3 in the context of (C3-SUMO1)_4_ and compared the results with C3 unfolding in the context of (C3-L)_4,_ both in TFP and OFP (Figure 3C,D, Table). Although the *mF_u_* of C3 in (C3-L)_4_ appears to be 12% higher in TFP experiments (*mF_u_* = 79.2 and 88.4 pN for the SUMO1- and L-containing heteropolyproteins, respectively), differences vanish in OFP experiments (*mF_u_* = 89.5 and 90.0 pN for the SUMO1- and L-containing heteropolyproteins, respectively). The RSD of the distribution of *∆mF_u_*, as estimated from Monte Carlo simulations fed with the actual number of experiments and events, is 2-3 times smaller in OFP than in TFP (Figure 3C,D, Table, Supplementary Figure 7B)

**Figure 3.**
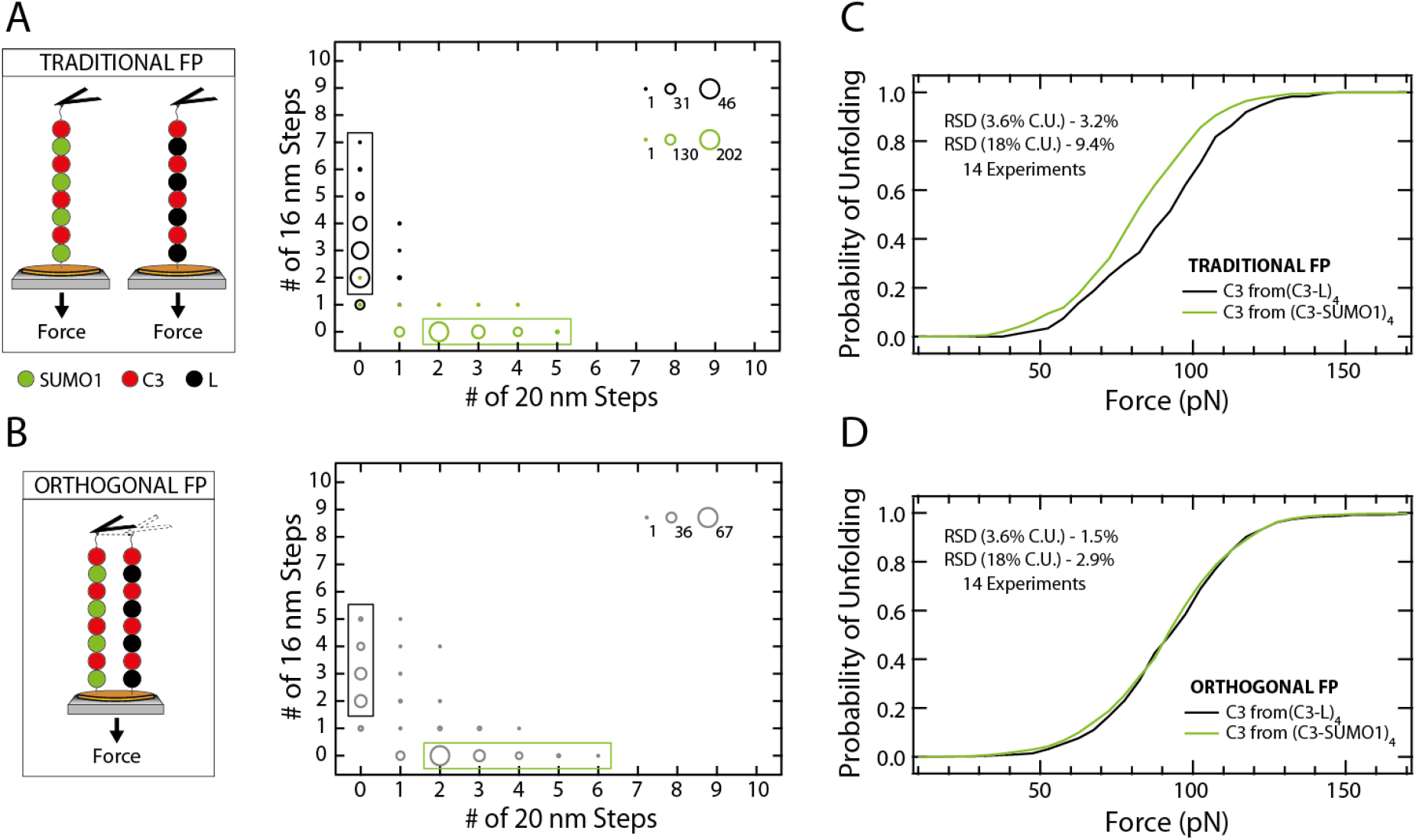
Dual-marker orthogonal fingerprinting overcomes confounding protein dimerization in concurrent atomic force spectroscopy. **(A)** Mechanical unfolding of (C3-SUMO1)_4_ and (C3-L)_4_ are measured in separate experiments by TFP. Individual traces are classified according to their number of 16 and 20 nm unfolding events, which mark the mechanical unfolding of protein L and SUMO1, respectively. The plot shows the frequency of the traces that have different combination of unfolding events, as indicated by the size of the dots. Traces coming from mechanical unfolding of (C3-SUMO1)_4_ are assigned when n(16nm) = 0 and n(20nm) > 1 (green rectangle), whereas (C3-L)_4_ traces are identified by n(16nm) > 1 and n(20nm) = 0. Number of traces analyzed: (C3-L)_4_, 163; (C3-SUMO1)_4_, 525. **(B)** Mechanical unfolding of (C3-SUMO1)_4_ and (C3-L)_4_, as measured concurrently in OFP experiments. The plot shows the frequency of the traces that have different combination of unfolding events, as indicated by the size of the dots (298 traces). The gating strategy defined in panel A allows the classification of the traces as originating from (C3-SUMO1)_4_ (green rectangle) or (C3-L)_4_ (black rectangle). **(C)** Experimental cumulative unfolding probability distribution of the C3 domain in the context of (C3-L)_4_ (6 experiments, 177 events, black; data is also presented in Figure 2C) and (C3-SUMO1)_4_ (8 experiments, 742 events, green), as measured in TFP. **(D)** Experimental cumulative unfolding probability distribution of the C3 domain in the context of (C3-L)_4_ (873 events, black) and (C3-SUMO1)_4_ (1043 events, green), resulting from unfolding data obtained in 14 independent OFP experiments. RSD values in the insets of panels C and D are estimated using Monte Carlo simulations that consider extreme values of calibration uncertainty (C.U.) (see also Supplementary Figure 7B). The pulling rate in all experiments was 40 pN/s.

### An error propagation model to further improve performance of concurrent atomic force spectroscopy

Our Monte Carlo simulations show that the improvement in accuracy in *∆mF_u_* of concurrent atomic force spectroscopy is preserved when multiple AFM experiments are averaged, and is independent of calibration uncertainty (Figure 1E,F). In apparent contradiction, we found that the RSDs of the distributions of *∆mF_u_* calculated for OFP concurrent datasets increase with calibration uncertainty (Table). Here, we propose a model for the propagation of calibration errors in concurrent single-molecule force-spectroscopy AFM experiments that accounts for all those observations. Considering that every unfolding force measured in the AFM is affected by an error δ as a consequence of inaccurate force calibration, the following equation for the measured *∆mF_u_* can be derived (Supplementary Text 3):

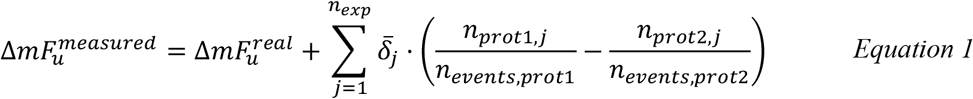

In Equation 1, 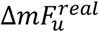 is the value of *∆mF_u_* that would be measured if there was no error in calibration, *n_exp_* is the number of experiments, *n_prot1,j_* and *n_prot2,j_* are the number of unfolding events for both proteins being compared in concurrent experiment *j*, *n_events,prot1_* and *n_events,prot2_* are the total number of unfolding events for each protein considering all experiments, and *δ̄_j_* is the average value of the error in force in experiment *j*, which is considered to be equivalent for two proteins measured concurrently.

Equation 1 shows that in concurrent measurements, 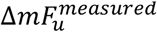 can differ from 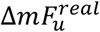 according to the magnitude of *δ̄_j_* values, which originate from the uncertainty in calibration, and to the specific number of unfolding events obtained for each protein in the different experiments contributing to the dataset. Importantly, if the proportion of events for both proteins is constant in all experiments (“balanced” condition: *n_prot1,j_* = *constant · n_prot2,j_*), Equation 1 leads to 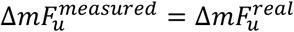, i.e. the relative mechanical stability measured in concurrent experiments is not affected by errors in calibration, independently of the number of experiments contributing to the dataset. Monte Carlo simulations readily confirm this prediction. While simulations of 2 concurrent experiments with 100 events per protein show that the resulting RSD of the distribution of *∆mF_u_* does not depend on the uncertainty in calibration, breaking the balanced condition by considering *n_prot_*_1_*,_exp_*_1_ = *n_prot_*_2_*,_exp_*_2_ = *150* and *n_prot_*_1_*, _exp_*_2_ = *n_prot_*_2*,exp*1_ = *50*, results in a dramatic increase of RSD especially at higher calibration uncertainties (Figure 4A). Under these unbalanced conditions, the performance of the concurrent strategy diminishes drastically and the obtained RSD approaches the one obtained in traditional atomic force spectroscopy (Figure 4A).

**Figure 4.**
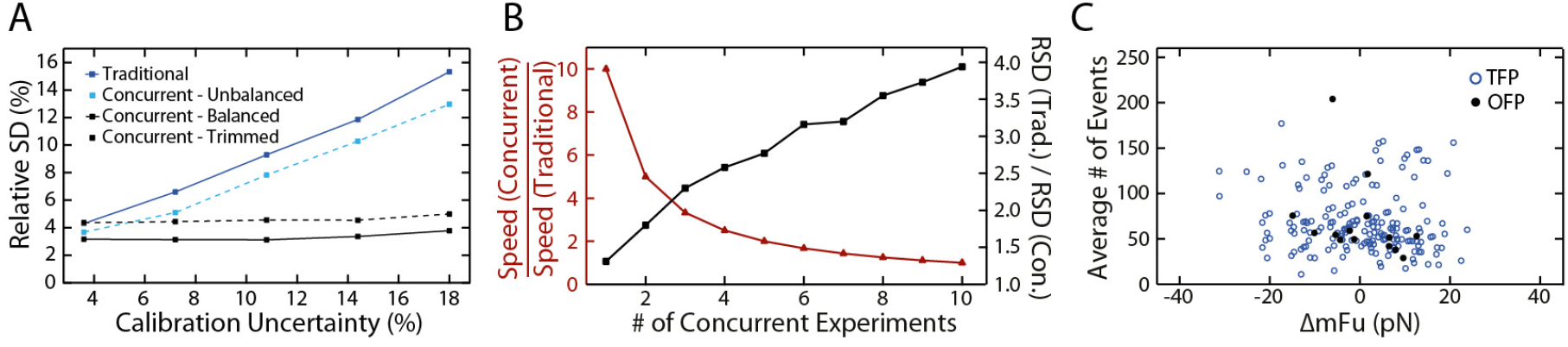
Improved accuracy and throughput of concurrent atomic force spectroscopy. **(A)** Monte Carlo simulations show that balancing datasets obtained in concurrent experiments improves RSD of distributions of *∆mF_u_* at calibration uncertainties >6%. Simulations of balanced datasets (solid lines, 100 events per experiment and protein) considered 2 experiments per protein in traditional (blue), and 2 experiments in concurrent AFM (black). To simulate unbalanced datasets, uneven number of unfolding events (50/150) were considered for both proteins in the first simulated experiment, and the order was reversed in the second simulated experiment (dashed blue line). The effects of balancing datasets (“trimming”) was examined by running simulations at the lowest number of events for both experiments (50, dashed black line). **(B)** Improvement in the speed of data acquisition (red) and RSD in the distribution of *∆mF_u_* (black) by concurrent measurements are estimated using Monte Carlo simulations at 10.8% calibration uncertainty (100 events per experiment and protein). Values are compared to a situation where two proteins are measured in 5 traditional experiments per protein. **(C)** The experimental *mF_u_* of C3 in the context of (C3-L)_4_ and (C3-SUMO1)_4_ is compared in 14 individual OFP experiments, and the corresponding *∆mF_u_* is represented (solid black circles). Using the same dataset, the equivalent *∆mF_u_* obtained in TFP was calculated from pairwise comparisons of *mF_u_* values from different experiments that have different force calibrations (open blue circles).

Since unbalanced data distributions result in poorer performance of concurrent atomic force spectroscopy, we examined whether balancing datasets through data removal could result in improved accuracy in *∆mF_u_*. We did Monte Carlo simulations of 2 concurrent experiments in which *n_prot1,j_* = *n_prot2,j_* = 50, i.e. we trimmed 100 extra events per protein so that both experiments had the same number of unfolding events for both proteins. As expected, the RSD of the distribution of *∆mF_u_* after trimming becomes independent of the calibration uncertainty (Figure 4A). Even though having less events per experiment results *per se* in poorer definition of distributions of unfolding forces (Figure 1C, Supplementary Figure 5B), the RSDs of the distributions of *∆mF_u_* obtained using trimmed datasets become lower than in the case of the more populated, but unbalanced, dataset at calibration uncertainties higher than 6% (Figure 4A).

Concurrent single-molecule atomic force spectroscopy datasets are expected to be unbalanced, since the different proteins are picked up randomly by the AFM cantilever (Supplementary Table 1). We have tested whether balancing our experimental OFP datasets also leads to improved accuracy in *∆mF_u_*. To this end, we have removed unfolding events so that every OFP experiment verifies the balanced condition *n_prot_*_1_ = *n_prot_*_2_. Using Monte Carlo simulations, we estimate that the RSD of the distribution of *∆mF_u_* obtained from the trimmed OFP datasets becomes lower than the original RSD also at calibration uncertainties higher than 6-7% (Supplementary Figure 7). In the two OFP datasets analyzed, the differences between the balanced and the unbalanced conditions are less prominent than in the case of artificial datasets consisting of only two experiments (Figure 4A). Indeed, we find that the extent of improvement in performance by dataset trimming decreases with the number of experiments (Supplementary Figure 9). Hence, we recommend that improvement in performance by balancing datasets is estimated on a case-by-case basis using Monte Carlo simulations fed with actual experimental data, as we have done here.

## DISCUSSION

In this report, we address a main limitation of force-spectroscopy by AFM that arises from uncertain calibration of cantilevers. We show, for the first time to our knowledge, that concurrent measurements result in remarkable improvements in the determination of relative mechanical properties by atomic force spectroscopy. We propose that concurrent AFM measurements of relative mechanical properties of cells, materials and tissues can result in gains in performance similar to those described here for proteins. Remarkably, concurrent strategies can be achieved by adapting already available atomic force spectroscopy techniques based on fluorescent labelling of cells or polymer blends ^31,32^.

Our work demonstrates that concurrent single-molecule atomic force spectroscopy outperforms traditional methods in three key aspects (Figure 5):

i. **The accuracy in *∆mF_u_* achieved by concurrent atomic force spectroscopy measurements is independent of calibration uncertainty even when multiple experiments are averaged** (Figures 1F,4A, Supplementary Figure 6). Although the uncertainty of cantilever calibration is usually considered to lie in the range of 5-25% depending on the calibration method ^15^, real uncertainties are extremely challenging to estimate. We have provided an experimental lower bound of 3.6% uncertainty in cantilever calibration by the commonly-used thermal fluctuations method (Supplementary Text 2). Uncertainties in spring constant calibration by the thermal fluctuations method have also been estimated by interlaboratory experiments, finding a value of up to 11% ^33^. However, neither of these approaches addresses more fundamental assumptions of the calibration procedures, or difficult-to-detect defects in individual cantilevers, which lead to higher calibration uncertainties ^34^. Indeed, we propose that concurrent atomic force spectroscopy strategies, in combination with further theoretical developments, can be key to exploit the advantages of next generation cantilevers, which are pushing the AFM into ranges of force, speed, stability and time resolution at the expense of more challenging force calibration ^14, 35-39^. It is worth mentioning that in concurrent strategies where the proteins are randomly picked up and mechanically probed, changes in *k_sc_* during experimentation affect the different proteins equally leading to preserved high accuracy in *∆mF_u_*.
ii. **Concurrent atomic force spectroscopy shows much improved accuracy in *∆mF_u_*** (Figures 1E,F, 4A,B, 5). Keeping the speed of data acquisition constant at high calibration uncertainties, the accuracy achieved by concurrent measurements can be 6 times higher than in traditional atomic force spectroscopy (Supplementary Figure 6B).
iii. **Concurrent atomic force spectroscopy leads to high-throughput relative mechanical characterization of proteins** (Figure 5). We estimate that concurrent atomic force spectroscopy can obtain the data needed for the same accuracy in *∆mF_u_* over 30 times faster than traditional measurements at high calibration uncertainties (Supplementary Figure 6C).

**Figure 5.**
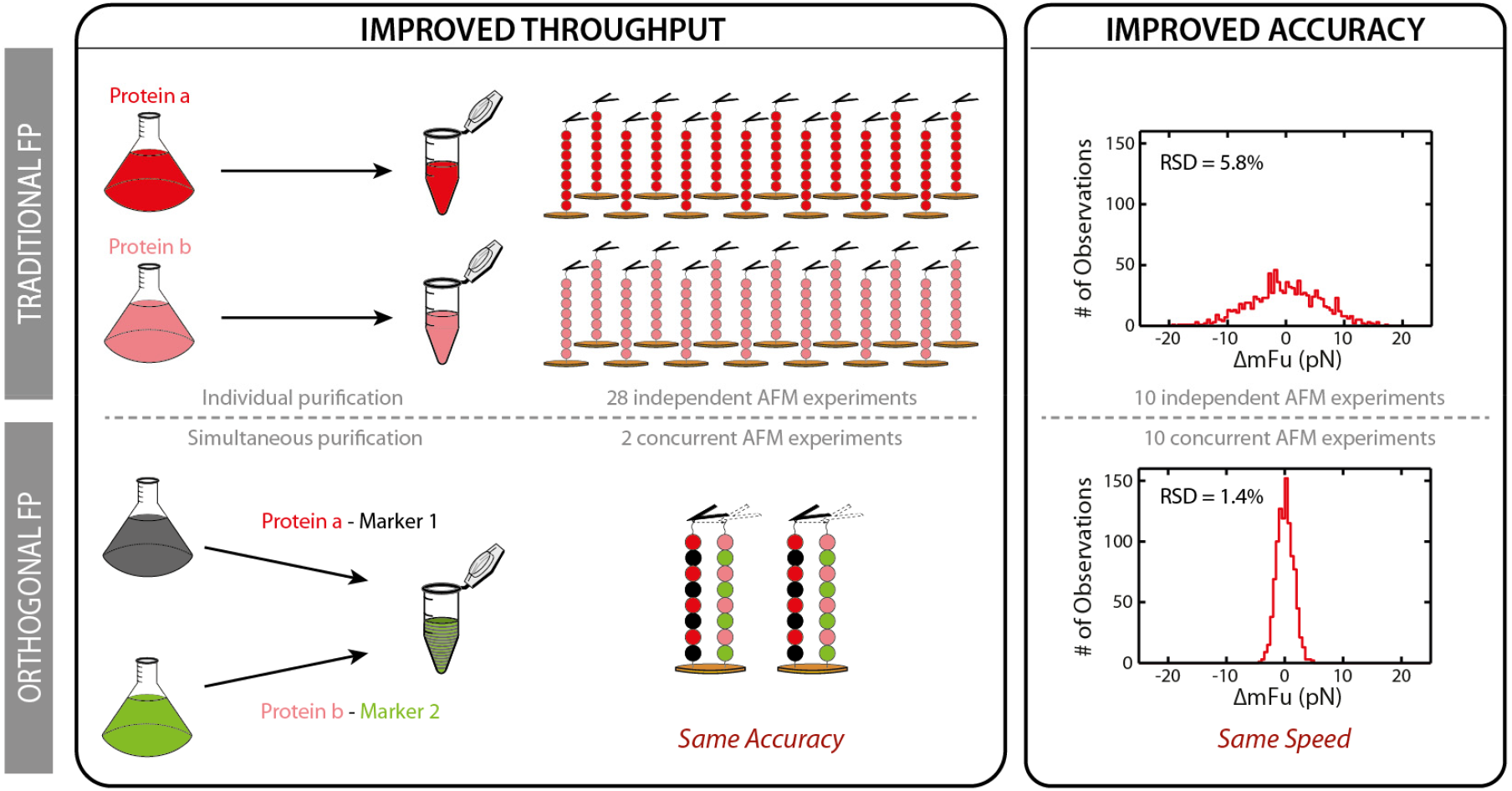
Orthogonal-fingerprinting-based concurrent atomic force spectroscopy. In the traditional approach, comparison of the mechanical stability of *Protein a* and *Protein b* involves independent purification and several AFM experiments to compensate for inaccurate force calibration. OFP is based on the production of heteropolyproteins composed of the proteins of interest fused to marker domains. Since the markers provide unequivocal fingerprints in single-molecule pulling experiments, OFP enables simultaneous purification and concurrent mechanical measurements, circumventing errors in force calibration. Concurrent measurements can achieve the same accuracy in *∆mF_u_* as conventional single-molecule atomic force spectroscopy with much better throughput (*left*). Alternatively, by keeping the speed of data acquisition constant, concurrent measurements considerably improve the accuracy in the determination of *∆mF_u_* (*right*). Improvements in throughput and accuracy are estimated from Monte Carlo simulations at 10.8% calibration uncertainty (100 events per experiment and protein).

The increase in throughput and accuracy in *∆mF_u_* of concurrent atomic force spectroscopy come at the expense of each other. Hence, depending on the goals of a particular study, the experimenter can choose to favor one or the other, or to find a balance between both. For instance, in Figure 4B, we show that different gains in accuracy and throughput can be achieved depending on the number of concurrent experiments carried out to compare unfolding forces of two proteins.

Motivated by our findings, we have developed OFP strategies for concurrent nanomechanical profiling of proteins. The increase in accuracy in *∆mF_u_* examined by Monte Carlo simulations is also captured in our limited experimental dataset, since the spread of *∆mF_u_* in pairs of OFP experiments is lower than in traditional experiments (SD = 8.0 *vs* 11.2 pN, respectively) (Figure 4C). An advantage of OFP over a previous concurrent strategy based on spatial separation of proteins ^22,23^ is that it can be readily implemented in any atomic force spectroscopy setup. In addition, different fingerprinting lengths in OFP provide additional reassurance of the identity of the probed proteins. In this regard, OFP is very well suited to compare mechanical properties of proteins with similar unfolding lengths. Hence, OFP can define mechanical hierarchies in multidomain proteins with higher accuracy, and lead to better descriptions of the mechanical effects of mutations, posttranslational modifications and the chemical environment on proteins and their complexes ^40-45^. Indeed, our highly accurate OFP experiments show that the mechanical stabilities of the C3 domain in the context of a (C3)_8_ homopolyprotein, or within a (C3-L)_4_ or (C3-SUMO1)_4_, are very similar (Figures 2E, 3D, Table). The fact that the mechanical stability of C3 does not depend on the neighboring marker domains lends strong support to the use of heteropolyproteins in force-spectroscopy AFM ^28, 46-48^. Another advantage of OFP is that orthogonally fingerprinted proteins can be purified simultaneously (Supplementary Figures 2, 10), resulting in extra savings in working time and reagents and ensuring equal experimental conditions for both proteins (Figure 5). Taking into account the full experimental workflow of single-molecule atomic force spectroscopy experiments, we estimate that the increase in throughput described in Figure 5 results in ~4 times less total experimentation time (Supplementary Figure 11).

We envision two features of OFP that can be further optimized. Since the relative performance of OFP is better at high number of events per experiment (Figure 1E, Supplementary Text 4), we propose that better improvements can be achieved by taking advantage of highly efficient high-strength single-molecule tethering methods ^23,29,49^. In addition, OFP strategies hold the promise of further parallelization by the use of multiple marker proteins. Importantly, in those cases where proteins have different unfolding lengths, concurrent single-molecule AFM measurements are of immediate application leading to the increase in accuracy and throughput described here ^21^. Examples that could benefit from application of concurrent measurements include examination of the effect of disulfide bonds, protein misfolding, multimerization, and pulling geometry in the mechanical stability of proteins and their complexes ^50-57^, and determination of rates of force-activated chemical reactions ^58^. In these examples, our Monte Carlo simulations can be applied to guide experimental design and interpretation (code is provided as Supplementary Material).

## METHODS

### Monte Carlo simulations

Monte Carlo simulations were programmed in Igor 6 (Wavemetrics). Simulations randomly assign an error in force according to a set uncertainty in the calibration between 3.6% and 18% (Supplementary Text 2). Every simulated atomic force spectroscopy experiment is therefore affected by a different calibration error, except under the condition of concurrent measurements, in which two proteins are probed in the same experiment and share error in force. Each cycle of the simulations returns a value of *∆mF_u_* for two proteins using Gaussian fits to their distribution of unfolding forces, obtained from a given number of independent experiments and unfolding events. We used a bin size of 25 pN when simulating artificial datasets, or 5 pN when feeding simulations with real datasets for better comparison with experimental distributions. Simulations calculate the RSD of the *∆mF_u_* distribution obtained from 1,000 cycles of the procedure. RSD is defined as the SD of the distribution of *∆mF_u_*, normalized by the theoretical value of *mF_u_* ^24^.

Distributions of unfolding forces are calculated through a kinetic Monte Carlo routine that considers that protein unfolding is exponentially dependent on force according to the Bell’s model ^59^:

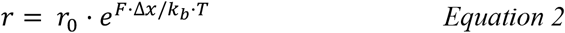

In Equation 2, *r* is the rate of protein unfolding, *r_0_* is the rate of unfolding at zero force, *F* is the force experienced by the protein, *∆x* is the distance to the transition state, *k_b_* is the Boltzmann constant and *T* is the absolute temperature. In the simulations, we considered that *k_b_ ·T* = 4.11 pN·nm, *r_0_* = 0.01 s^-1^and *∆x* = 0.2 nm. We chose these values because they are a good estimate of mechanical unfolding parameters of C3. We checked that values of RSD are fairly insensitive to small variations in *r_0_* and *∆x* and therefore one single set of parameters is enough to calculate RSD of distributions of *∆mF_u_* even if the mechanical parameters of the proteins to be compared are slightly different (Supplementary Table 2).

The kinetic Monte Carlo to obtain distribution of unfolding forces compares a random number with the instantaneous probability of unfolding at a given force. If the unfolding probability is higher than the random number, unfolding is considered to happen at that force. Instantaneous probabilities of unfolding are calculated following a linear approximation ^60^:

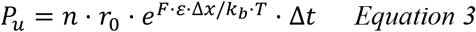

In Equation 3, *n* is the number of domains that remain folded at a particular force, *E* is the error in force due to the uncertain cantilever calibration (Supplementary Text 2) and *∆t* is the time step of the Monte Carlo. In the simulations, we used *∆t* = 10^−4^ s, which ensures validity of the linear approximation, since *n · r · ∆t* is kept under 0.05 (Supplementary Text 5). Pilot simulations show that results do not vary if we use a smaller time step of 10^−5^ s.

### Protein production and purification

The cDNAs coding for the C3-L and C3-SUMO1 constructs were produced by gene synthesis (NZY-Tech and Gene Art, respectively). The cDNA coding for the C3 domain was obtained by PCR. cDNAs coding for polyproteins were produced following an iterative strategy of cloning using BamHI, BglII and KpnI restriction enzymes, as described before ^27,61^. Final cDNAs were inserted in the pQE80L expression plasmid using BamHI and KpnI and the resulting plasmids were verified by Sanger sequencing. Full protein sequences are reported in Supplementary Text 1. Polyproteins were expressed in BLR (DE3) *E. coli* strain. Briefly, fresh cultures (OD_600_ = 0.6- 1.0) are induced with 1mM IPTG for 3 hours at 37°C and at 250 rpm. Cells are lysed by a combination of tip sonication and passes through a French Press. Polyproteins are purified from the soluble fraction through Ni-NTA agarose chromatography (Qiagen), following the recommendations of the supplier and adding 10 mM DTT to the buffers. Further purification is achieved by size-exclusion chromatography in an AKTA Pure 25L system using a Superose 6 Increase 10/300 GL or a Superdex 200 Increase 10/300 GL column (GE Healthcare). The proteins are eluted in 10 mM Hepes, pH 7.2, 150 mM NaCl, 1 mM EDTA, which is also the buffer used in atomic force spectroscopy experiments. Purity of samples was evaluated using SDS-PAGE gels (a typical result is shown in Supplementary Figure 2).

### Force-spectroscopy by AFM

Single-molecule AFM measurements were obtained in an AFS setup (Luigs & Neumann) according to established protocols ^7^. We used silicon nitride MLCT-C cantilevers with a 60-nm gold back side coating (Bruker), which we calibrated according to the thermal fluctuations method ^62^. Typical spring constant values ranged between 15 and 20 pN/nm. To run single-molecule AFM experiments, a small aliquot (2-10 μL) of the purified protein is deposited on the surface of a gold coated cover slip (Luigs & Neumann), or directly into the Hepes buffer contained in the fluid chamber of the AFS. The cantilever is brought in contact to the surface for 1-2 s at 500-2000 pN to favor formation of single-molecule tethers. Then, the surface is retracted to achieve the set point force. If a single-molecule tether is formed, the force is increased linearly at 40 pN/s for 5 s while the length of the polyprotein is measured. This protocol ensures full unfolding of C3, L and SUMO1 domains (Supplementary Figure 3). Unfolding events are detected as increases in the length of the protein. In the initial characterization of polyproteins, we analyze all traces that contain at least two events of the same size, which allows to set a fingerprinting length for the domains (24 ± 1 nm for C3, 16 ± 1 nm for protein L, and 20 ± 1 nm for SUMO1, see Supplementary Figure 3). For the rest of the analyses, we only considered traces that contain fingerprinting unfolding lengths. Unfolding forces were recorded and plotted as cumulative distributions. *mF_u_* values were obtained from Gaussian fits to histograms of unfolding forces. Force inaccuracy due to laser interference was lower than 40 pN in all experiments (peak-to-peak height in baseline force-extension recordings) ^7^.

## Author contributions

J.A.C. designed the research. D.V.C engineered polyprotein constructs and produced proteins. C.P.L., C.S.C and D.S.O. did AFM experiments and analyzed single-molecule data. J.A.C programmed Monte Carlo simulations. C.P.L., C.S.C and J.A.C. run and analyzed Monte Carlo simulations and assembled display figures. J.A.C. drafted the manuscript with input from all the authors.

## Acknowledgements

J.A.C. acknowledges funding from the Spanish Ministry of Economy, Industry and Competitiveness (MEIC) through grants BIO2014-54768-P, BIO2017-83640-P, and RYC-2014-16604, the European Research Area Network on Cardiovascular Diseases (ERA-CVD, MINOTAUR, AC16/00045), and the CNIC intramural grant program (03-2016 IGP). The CNIC is supported by MEIC and the Pro CNIC Foundation, and is a Severo Ochoa Center of Excellence (SEV-2015-0505). C.P.L. was a recipient of CNIC Master Fellowship. C.S.C. is the recipient of an FPI predoctoral fellowship BES-2016-076638. We thank Natalia Vicente for excellent technical support. We thank all the members of the laboratory of Molecular Mechanics of the Cardiovascular System for helpful discussions. We thank Elías Herrero-Galán (CNIC, Madrid) for critical reading of the manuscript and Lidia Prieto-Frías for key insights about propagation of calibration errors.

## Resources

The code used for the Monte Carlo simulations is available as online Supplementary Material.

## REFERENCES

1. Binnig, G., Quate, C. F. & Gerber, C. Atomic force microscope. Physical review letters 56, 930–933. (1986).

2. Garcia, R. & Herruzo, E. T. The emergence of multifrequency force microscopy. Nature nanotechnology 7, 217–226. (2012).

3. Krieg, M., Fläschner, G., Alsteens, D., Gaub, B. M., Roos, W. H., Wuite, G. J. L., Gaub, H. E., Gerber, C., Dufrêne, Y. F. & Müller, D. J. Atomic force microscopy-based mechanobiology. Nature Reviews Physics(2018).

4. Ando, T., Uchihashi, T. & Kodera, N. High-Speed AFM and Applications to Biomolecular Systems. Annual Review of Biophysics 42, 393–414. (2013).

5. Al-Rekabi, Z. & Contera, S. Multifrequency AFM reveals lipid membrane mechanical properties and the effect of cholesterol in modulating viscoelasticity. Proceedings of the National Academy of Sciences(2018).

6. Lai, C.-Y., Santos, S. & Chiesa, M. Systematic Multidimensional Quantification of Nanoscale Systems From Bimodal Atomic Force Microscopy Data. ACS Nano 10, 6265–6272. (2016).

7. Popa, I., Kosuri, P., Alegre-Cebollada, J., Garcia-Manyes, S. & Fernandez, J. M. Force dependency of biochemical reactions measured by single-molecule force-clamp spectroscopy. Nat Protoc 8, 1261–1276. (2013).

8. Neuman, K. C. & Nagy, A. Single-molecule force spectroscopy: optical tweezers, magnetic tweezers and atomic force microscopy. Nat Methods 5, 491–505. (2008).

9. Rief, M., Gautel, M., Oesterhelt, F., Fernandez, J. M. & Gaub, H. E. Reversible unfolding of individual titin immunoglobulin domains by AFM. Science 276, 1109–1112. (1997).

10. Linke, W. A. & Hamdani, N. Gigantic business: titin properties and function through thick and thin. Circ Res 114, 1052–1068. (2014).

11. del Rio, A., Perez-Jimenez, R., Liu, R., Roca-Cusachs, P., Fernandez, J. M. & Sheetz, M. P. Stretching single talin rod molecules activates vinculin binding. Science 323, 638–641. (2009).

12. Paluch, E. K., Nelson, C. M., Biais, N., Fabry, B., Moeller, J., Pruitt, B. L., Wollnik, C., Kudryasheva, G., Rehfeldt, F. & Federle, W. Mechanotransduction: use the force(s). BMC biology 13, 47. (2015).

13. Li, H., Linke, W. A., Oberhauser, A. F., Carrion-Vazquez, M., Kerkvliet, J. G., Lu, H., Marszalek, P. E. & Fernandez, J. M. Reverse engineering of the giant muscle protein titin. Nature 418, 998–1002. (2002).

14. Slattery, A. D., Blanch, A. J., Ejov, V., Quinton, J. S. & Gibson, C. T. Spring constant calibration techniques for next-generation fast-scanning atomic force microscope cantilevers. Nanotechnology 25, 335705. (2014).

15. Charles, A. C. & Martin, P. S. The determination of atomic force microscope cantilever spring constants via dimensional methods for nanomechanical analysis. Nanotechnology 16, 1666. (2005).

16. Killgore, J. P., Geiss, R. H. & Hurley, D. C. Continuous measurement of atomic force microscope tip wear by contact resonance force microscopy. Small 7, 1018–1022. (2011).

17. Wagner, R., Moon, R., Pratt, J., Shaw, G. & Raman, A. Uncertainty quantification in nanomechanical measurements using the atomic force microscope. Nanotechnology 22, 455703. (2011).

18. Schillers, H., Rianna, C., Schape, J., Luque, T., Doschke, H., Walte, M., Uriarte, J. J., Campillo, N., Michanetzis, G. P. A., Bobrowska, J., Dumitru, A., Herruzo, E. T., Bovio, S., Parot, P., Galluzzi, M., Podesta, A., Puricelli, L., Scheuring, S., Missirlis, Y., Garcia, R., Odorico, M., Teulon, J. M., Lafont, F., Lekka, M., Rico, F., Rigato, A., Pellequer, J. L., Oberleithner, H., Navajas, D. & Radmacher, M. Standardized Nanomechanical Atomic Force Microscopy Procedure (SNAP) for Measuring Soft and Biological Samples. Scientific reports 7, 5117. (2017).

19. Anderson, B. R., Bogomolovas, J., Labeit, S. & Granzier, H. Single molecule force spectroscopy on titin implicates immunoglobulin domain stability as a cardiac disease mechanism. J Biol Chem 288, 5303–5315. (2013).

20. Sadler, D. P., Petrik, E., Taniguchi, Y., Pullen, J. R., Kawakami, M., Radford, S. E. & Brockwell, D. J. Identification of a mechanical rheostat in the hydrophobic core of protein L. J Mol Biol 393, 237–248. (2009).

21. Jobst, M. A., Milles, L. F., Schoeler, C., Ott, W., Fried, D. B., Bayer, E. A., Gaub, H. E. & Nash, M. A. Resolving dual binding conformations of cellulosome cohesin-dockerin complexes using single-molecule force spectroscopy. eLife 4(2015).

22. Otten, M., Ott, W., Jobst, M. A., Milles, L. F., Verdorfer, T., Pippig, D. A., Nash, M. A. & Gaub, H. E. From genes to protein mechanics on a chip. Nat Methods 11, 1127–1130. (2014).

23. Verdorfer, T., Bernardi, R. C., Meinhold, A., Ott, W., Luthey-Schulten, Z., Nash, M. A. & Gaub, H. E. Combining in Vitro and in Silico Single-Molecule Force Spectroscopy to Characterize and Tune Cellulosomal Scaffoldin Mechanics. Journal of the American Chemical Society 139, 17841–17852. (2017).

24. Schlierf, M., Li, H. & Fernandez, J. M. The unfolding kinetics of ubiquitin captured with single-molecule force-clamp techniques. Proc Natl Acad Sci U S A 101, 7299–7304. (2004).

25. Alegre-Cebollada, J., Kosuri, P., Giganti, D., Eckels, E., Rivas-Pardo, J. A., Hamdani, N., Warren, C. M., Solaro, R. J., Linke, W. A. & Fernandez, J. M. S-glutathionylation of cryptic cysteines enhances titin elasticity by blocking protein folding. Cell 156, 1235–1246. (2014).

26. Carrion-Vazquez, M., Oberhauser, A. F., Fisher, T. E., Marszalek, P. E., Li, H. & Fernandez, J. M. Mechanical design of proteins studied by single-molecule force spectroscopy and protein engineering. Prog Biophys Mol Biol 74, 63–91. (2000).

27. Carrion-Vazquez, M., Oberhauser, A. F., Fowler, S. B., Marszalek, P. E., Broedel, S. E., Clarke, J. & Fernandez, J. M. Mechanical and chemical unfolding of a single protein: a comparison. Proc Natl Acad Sci U S A 96, 3694–3699. (1999).

28. Li, H., Oberhauser, A. F., Redick, S. D., Carrion-Vazquez, M., Erickson, H. P. & Fernandez, J. M. Multiple conformations of PEVK proteins detected by single-molecule techniques. Proceedings of the National Academy of Sciences of the United States of America 98, 10682–10686. (2001).

29. Popa, I., Berkovich, R., Alegre-Cebollada, J., Badilla, C. L., Rivas-Pardo, J. A., Taniguchi, Y., Kawakami, M. & Fernandez, J. M. Nanomechanics of HaloTag tethers. Journal of the American Chemical Society 135, 12762–12771. (2013).

30. Kotamarthi, H. C., Sharma, R. & Koti Ainavarapu, S. R. Single-molecule studies on PolySUMO proteins reveal their mechanical flexibility. Biophysical journal 104, 2273–2281. (2013).

31. Herruzo, E. T., Perrino, A. P. & Garcia, R. Fast nanomechanical spectroscopy of soft matter. Nature communications 5, 3126. (2014).

32. Bhat, S. V., Sultana, T., Körnig, A., McGrath, S., Shahina, Z. & Dahms, T. E. S. Correlative atomic force microscopy quantitative imaging-laser scanning confocal microscopy quantifies the impact of stressors on live cells in real-time. Scientific reports 8, 8305. (2018).

33. te Riet, J., Katan, A. J., Rankl, C., Stahl, S. W., van Buul, A. M., Phang, I. Y., Gomez-Casado, A., Schon, P., Gerritsen, J. W., Cambi, A., Rowan, A. E., Vancso, G. J., Jonkheijm, P., Huskens, J., Oosterkamp, T. H., Gaub, H., Hinterdorfer, P., Figdor, C. G. & Speller, S. Interlaboratory round robin on cantilever calibration for AFM force spectroscopy. Ultramicroscopy 111, 1659–1669. (2011).

34. Ohler, B. Practical Advice on the Determination of Cantilever Spring Constants. Veeco Instruments Incorporated, Technical Report., (2007).

35. Rico, F., Gonzalez, L., Casuso, I., Puig-Vidal, M. & Scheuring, S. High-speed force spectroscopy unfolds titin at the velocity of molecular dynamics simulations. Science 342, 741–743. (2013).

36. Yu, H., Siewny, M. G., Edwards, D. T., Sanders, A. W. & Perkins, T. T. Hidden dynamics in the unfolding of individual bacteriorhodopsin proteins. Science 355, 945–950. (2017).

37. He, C., Hu, C., Hu, X., Hu, X., Xiao, A., Perkins, T. T. & Li, H. Direct Observation of the Reversible Two-State Unfolding and Refolding of an alpha/beta Protein by Single-Molecule Atomic Force Microscopy. Angewandte Chemie 54, 9921–9925. (2015).

38. Faulk, J. K., Edwards, D. T., Bull, M. S. & Perkins, T. T. Improved Force Spectroscopy Using Focused-Ion-Beam-Modified Cantilevers. Methods in enzymology 582, 321–351. (2017).

39. Uhlig, M. R., Amo, C. A. & Garcia, R. Dynamics of breaking intermolecular bonds in high-speed force spectroscopy. Nanoscale 10, 17112–17116. (2018).

40. Oroz, J., Bruix, M., Laurents, D. V., Galera-Prat, A., Schonfelder, J., Canada, F. J. & Carrion-Vazquez, M. The Y9P Variant of the Titin I27 Module: Structural Determinants of Its Revisited Nanomechanics. Structure 24, 606–616. (2016).

41. Li, H., Carrion-Vazquez, M., Oberhauser, A. F., Marszalek, P. E. & Fernandez, J. M. Point mutations alter the mechanical stability of immunoglobulin modules. Nat Struct Biol 7, 1117–1120. (2000).

42. Oberhauser, A. F., Badilla-Fernandez, C., Carrion-Vazquez, M. & Fernandez, J. M. The mechanical hierarchies of fibronectin observed with single-molecule AFM. J Mol Biol 319, 433–447. (2002).

43. Li, H., Oberhauser, A. F., Fowler, S. B., Clarke, J. & Fernandez, J. M. Atomic force microscopy reveals the mechanical design of a modular protein. Proc Natl Acad Sci U S A 97, 6527–6531. (2000).

44. Williams, P. M., Fowler, S. B., Best, R. B., Toca-Herrera, J. L., Scott, K. A., Steward, A. & Clarke, J. Hidden complexity in the mechanical properties of titin. Nature 422, 446–449. (2003).

45. Milles, L. F., Schulten, K., Gaub, H. E. & Bernardi, R. C. Molecular mechanism of extreme mechanostability in a pathogen adhesin. 359, 1527–1533. (2018).

46. Dietz, H. & Rief, M. Exploring the energy landscape of GFP by single-molecule mechanical experiments. Proceedings of the National Academy of Sciences of the United States of America 101, 16192–16197. (2004).

47. Steward, A., Toca-Herrera, J. L. & Clarke, J. Versatile cloning system for construction of multimeric proteins for use in atomic force microscopy. Protein Science 11, 2179–2183. (2002).

48. Oroz, J., Hervás, R. & Carrión-Vázquez, M. Unequivocal Single-Molecule Force Spectroscopy of Proteins by AFM Using pFS Vectors. Biophysical journal 102, 682–690. (2012).

49. Zakeri, B., Fierer, J. O., Celik, E., Chittock, E. C., Schwarz-Linek, U., Moy, V. T. & Howarth, M. Peptide tag forming a rapid covalent bond to a protein, through engineering a bacterial adhesin. Proceedings of the National Academy of Sciences of the United States of America 109, E690–697. (2012).

50. Giganti, D., Yan, K., Badilla, C. L., Fernandez, J. M. & Alegre-Cebollada, J. Disulfide isomerization reactions in titin immunoglobulin domains enable a mode of protein elasticity. Nature communications 9, 185. (2018).

51. Manteca, A., Schonfelder, J., Alonso-Caballero, A., Fertin, M. J., Barruetabena, N., Faria, B. F., Herrero-Galan, E., Alegre-Cebollada, J., De Sancho, D. & Perez-Jimenez, R. Mechanochemical evolution of the giant muscle protein titin as inferred from resurrected proteins. Nature structural & molecular biology 24, 652–657. (2017).

52. Manteca, A., Alonso-Caballero, A., Fertin, M., Poly, S., De Sancho, D. & Perez-Jimenez, R. The influence of disulfide bonds on the mechanical stability of proteins is context dependent. J Biol Chem 292, 13374–13380. (2017).

53. Bertz, M., Buchner, J. & Rief, M. Mechanical stability of the antibody domain CH3 homodimer in different oxidation states. J Am Chem Soc 135, 15085–15091. (2013).

54. Carrion-Vazquez, M., Li, H., Lu, H., Marszalek, P. E., Oberhauser, A. F. & Fernandez, J. M. The mechanical stability of ubiquitin is linkage dependent. Nat Struct Biol 10, 738–743. (2003).

55. Oberhauser, A. F., Marszalek, P. E., Carrion-Vazquez, M. & Fernandez, J. M. Single protein misfolding events captured by atomic force microscopy. Nat Struct Biol 6, 1025–1028. (1999).

56. Sarkar, A., Caamano, S. & Fernandez, J. M. The mechanical fingerprint of a parallel polyprotein dimer. Biophysical journal 92, L36–38. (2007).

57. Carl, P., Kwok, C. H., Manderson, G., Speicher, D. W. & Discher, D. E. Forced unfolding modulated by disulfide bonds in the Ig domains of a cell adhesion molecule. Proc Natl Acad Sci U S A 98, 1565–1570. (2001).

58. Garcia-Manyes, S., Liang, J., Szoszkiewicz, R., Kuo, T. L. & Fernandez, J. M. Force-activated reactivity switch in a bimolecular chemical reaction. Nature Chemistry 1, 236–242. (2009).

59. Bell, G. I. Models for the specific adhesion of cells to cells. Science 200, 618–627. (1978).

60. Shi, J., Gafni, A. & Steel, D. Simulated data sets for single molecule kinetics: some limitations and complications of data analysis. European biophysics journal: EBJ 35, 633–645. (2006).

61. Alegre-Cebollada, J., Badilla, C. L. & Fernandez, J. M. Isopeptide bonds block the mechanical extension of pili in pathogenic Streptococcus pyogenes. J Biol Chem 285, 11235–11242. (2010).

62. Hutter, J. L. & Bechhoefer, J. Calibration of atomic-force microscope tips. Review of Scientific Instruments 64, 1868–1873. (1993).

